## Supplementary material for "Encircling the regions of the pharmacogenomic landscape that determine drug response": Supp Info

### Additional file 1

#### Supplementary Figures

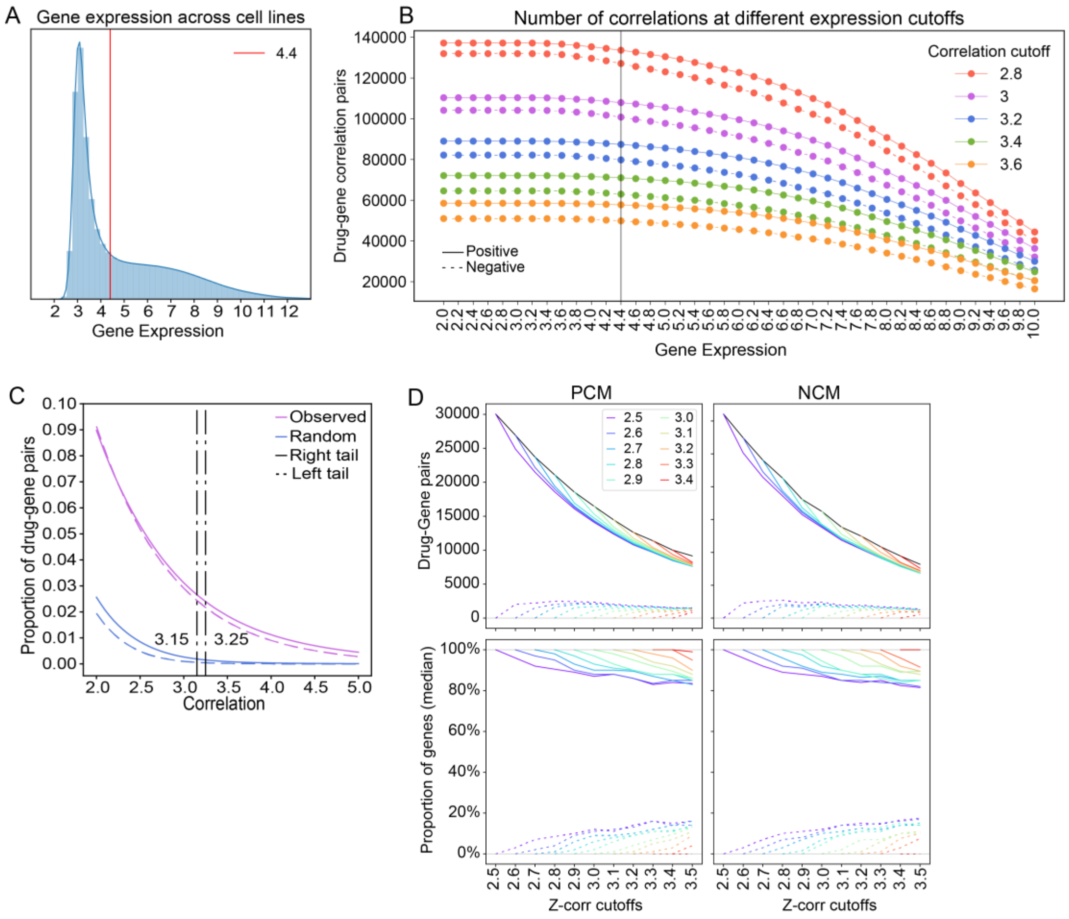


**Figure S1.** (A) Distribution of gene expression values across the GDSC cell line panel. The chosen cutoff of 4.4 is shown in red. (B) Robustness of the previous cutoff, measured as number of drug-gene correlations found. Continuous lines correspond to positive correlations and dashed lines to negative correlations. The colors of the lines denote different possible $z_{cor}$ values; the chosen one was 3.2 (in blue) (see next panel). (C) Absolute $z_{cor}$ for the observed and randomized gene-drug correlations. We chose a cutoff of 3.2, as it corresponded to well-accepted “p-values” of 0.05 and 0.001 in the observed and randomized distributions, respectively. (D) (Top panels) The black line denotes the number of drug-gene pairs encountered in modules (PCMs and NCMs) as a function of the $z_{cor}$ score cutoff [range 2.5-3.5]. The continuous colored lines show the number of drug-gene pairs “conserved” in the modules as we move to higher $z_{cor}$, with respect to the cutoff specified in the legend. On the contrary, dashed lines denote the genes that are added. (Bottom panels) Normalized version of the top panels, taking the total number of drug gene pairs (the black line) as a reference (100%).

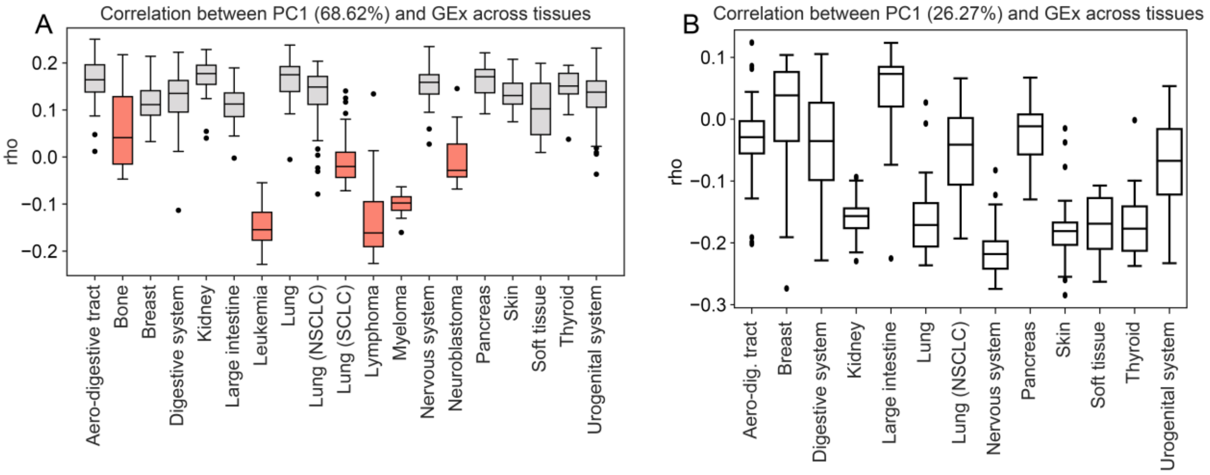


**Figure S2**. (A) We performed a Principal Component Analysis (PCA) in the drug-gene correlation distribution and kept the first principal component (PC1, explaining 65.63% of the variance). Then, we correlated the PC1 loadings to basal gene expression of each CCL (rho). In light of the results, we removed CCLs derived from neuroblastomas, hematopoietic, bone and small cell lung cancer tissues (in red) due to their characteristic rho values. (B) Analysis of the tissue effect in drug-gene correlations after filtering the most influential tissues. After the filtering, PC1 explains only 26% of the total variance and none of the tissues is distinctively correlated.

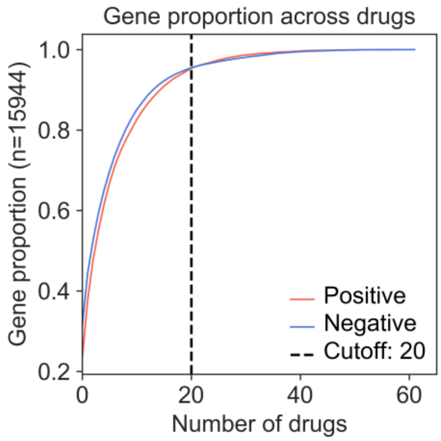


**Figure S3*.*** We counted the number times a positively (red) or negatively (blue) correlated gene appeared across drugs and plotted their cumulative distribution. The dashed black line shows the cutoff applied to identify frequently-correlated genes (5%).

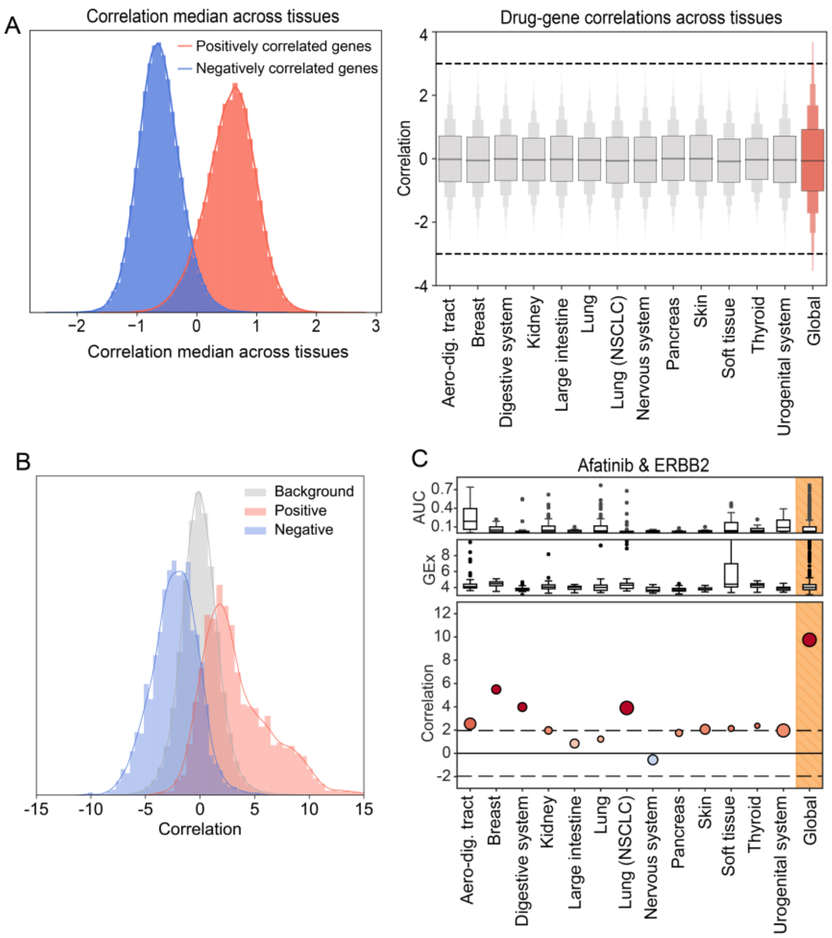


**Figure S4**. (A) (Left) Distribution of drug-gene correlation medians across tissues for positively (red) and negatively (blue) correlated genes ($z_{cor}$ beyond +/- 3.2). (Right) Drug-gene correlation distribution in each tissue. The right-most boxplot (in red) shows the correlations using all the tissues. (B) We calculated drug-gene correlation in an external dataset (CTRP panel) and identified positive and negative correlations (+/- 3.2 zcor). When mapped CTRP-correlation pairs on the GDSC results, CTRP-positive and CTRP-negative correlations were also found positively (red) and negatively (blue) correlated in GDSC, respectively. (C) Afatinib-ERBB2 correlation across tissues.

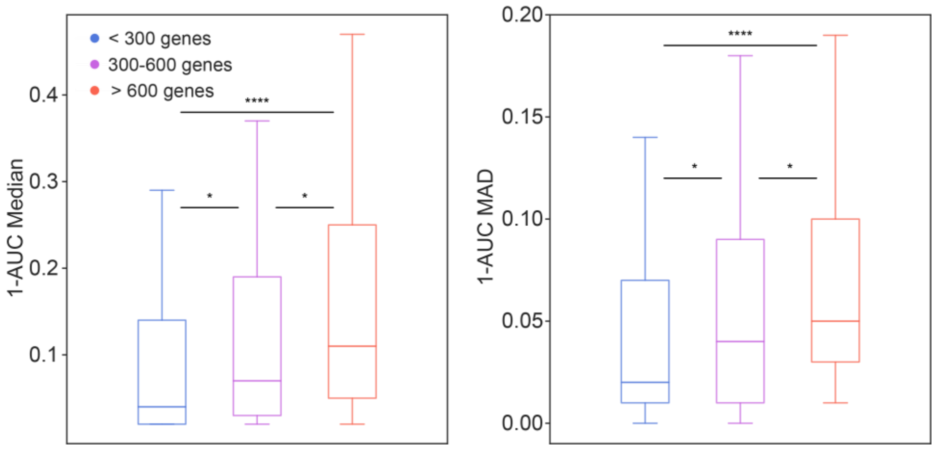


**Figure S5.** Median (left) and median absolute deviation (MAD, right) of 1-AUC values per drug across CCLs. Results are split by the number of genes (<300, 300-600 and >600) correlated to each drug. Globally, higher (left) and more variable (right) 1-AUC values allow for more drug-gene correlations to be detected.

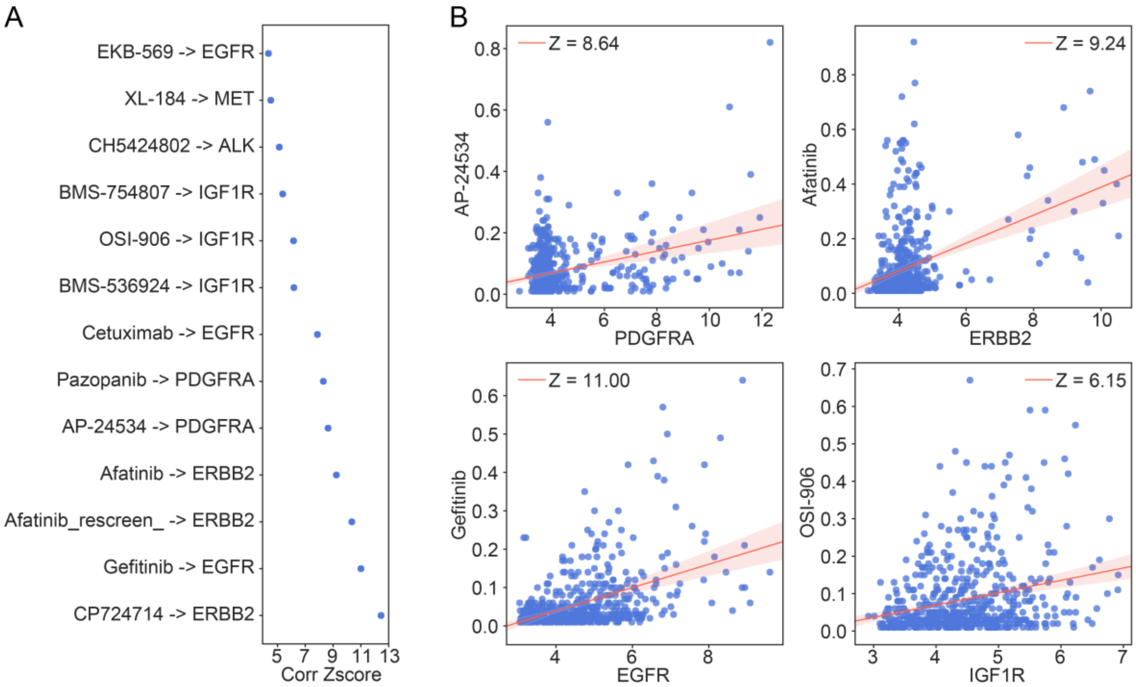


**Figure S6**. (A) Correlation between drugs and cell surface receptor targets. (B) Four exemplary drugs whose nominal target gene expression correlates to cell line sensitivity.

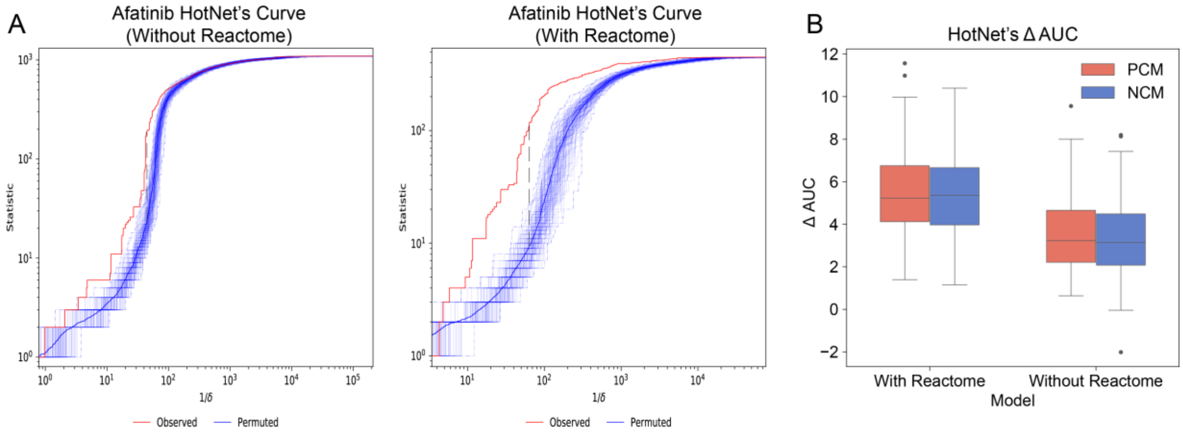


**Figure S7.** (A) HotNet statistic for the positively-correlated module (PCM) detection in Afatinib. Red lines correspond to observed data and blue lines to random runs. In the left panel, HotNet2 was run without pre-filtering. In the right panel, a Reactome-based pre-filtering was applied. (B) Difference between observed and random HotNet2 curves (quantified as the subtraction of areas) across all drugs, with and without the Reactome filtering.

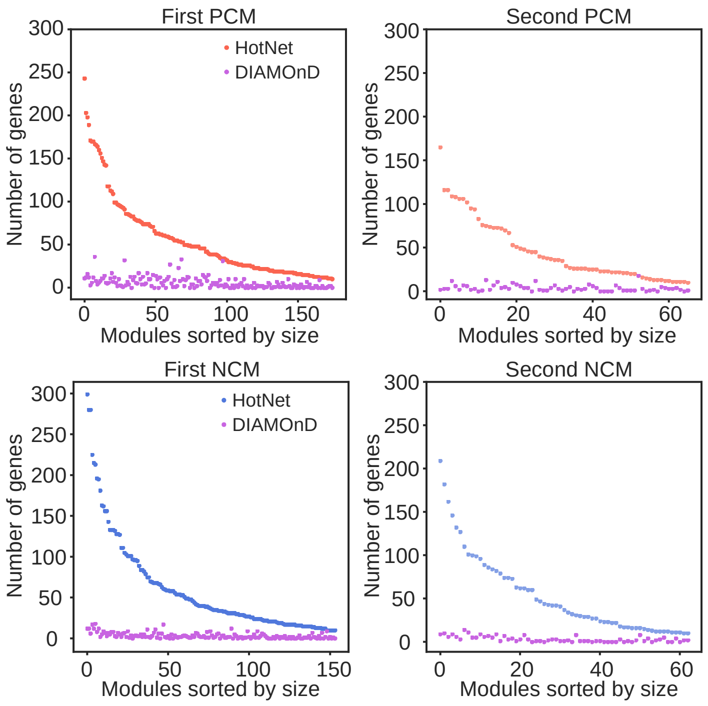


**Figure S8.** Number of genes added by the DIAMOND step in relation to genes added by HotNet2.

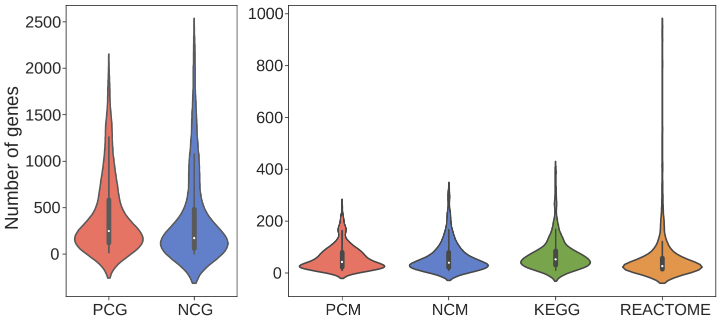


**Figure S9.** Number of positively and negatively correlated genes (PCGs, NCGs) per drug. Number of genes in positively and negatively correlated modules (PCMs, NCMs), compared to number of genes in KEGG and Reactome pathways.

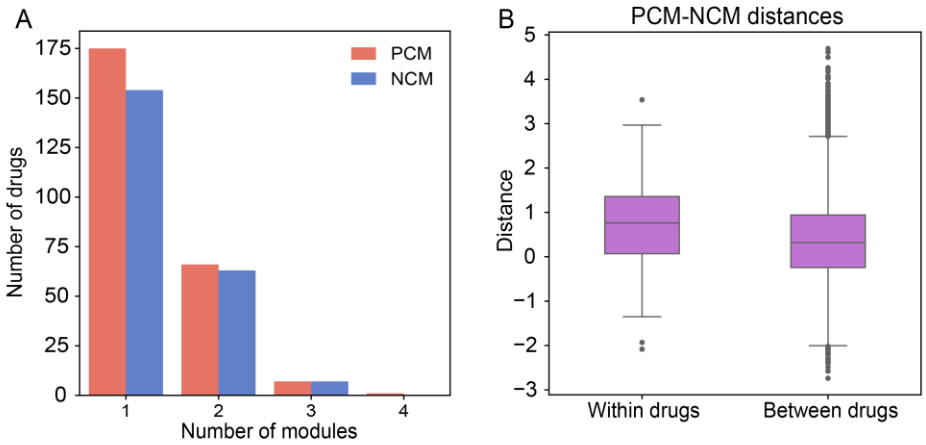


**Figure S10.** (A) Number of modules identified per drug (PCMs in red, NCMs in blue). (B) PCM-vs-NCM distances *within* drugs and *between* drugs.

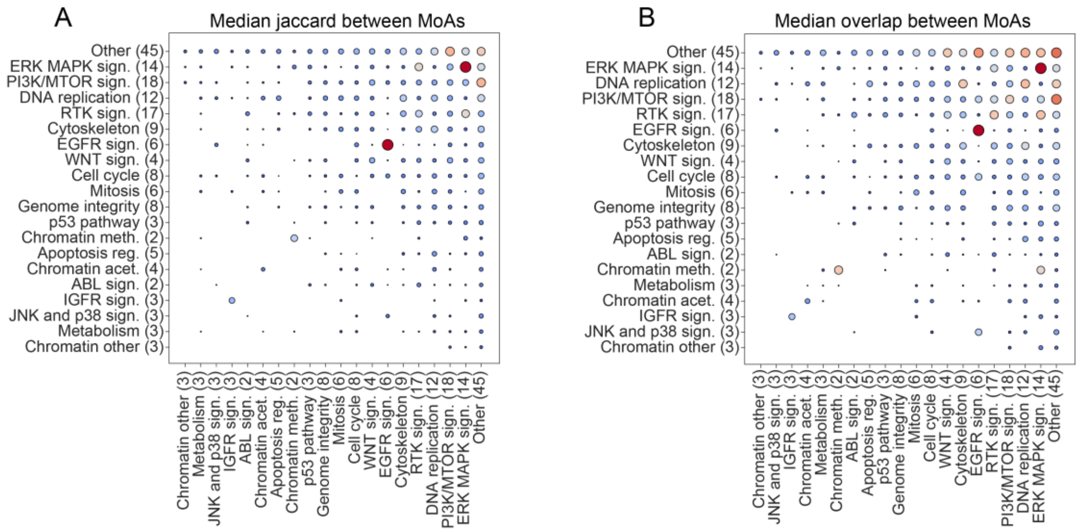


**Figure S11.** "Similarities" between drug modules of different drug classes. Larger and redder dots denote higher similarities. (A) Median Jaccard coefficient between genes [capped at 0.3 in the plot scale]. (B) Median overlap index (x-axis with respect to y-axis) [capped at 0.5].

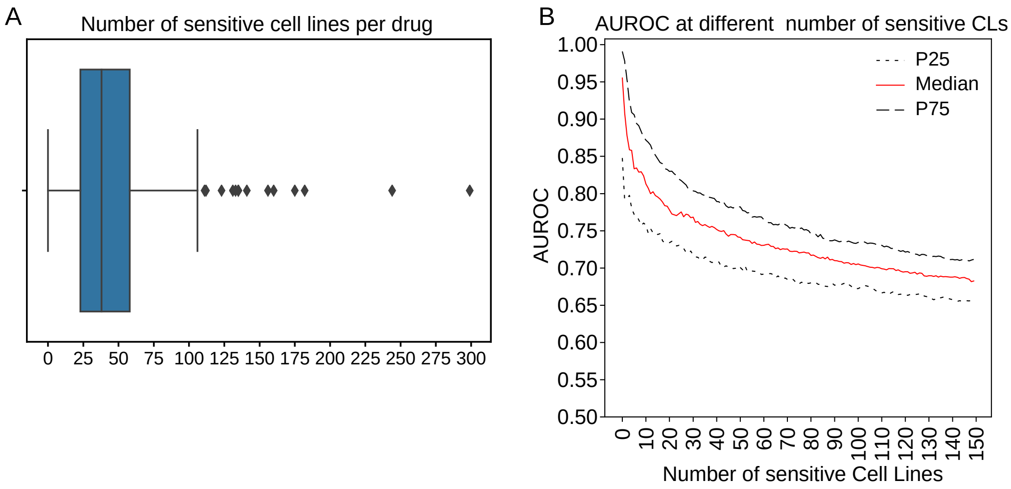


**Figure S12.** (A) Number of sensitive cell lines per drug according to the GDSC publication. (B) Predictive capability (AUROC) of the top-n sensitive cell lines, n ranging from 1 to 150.

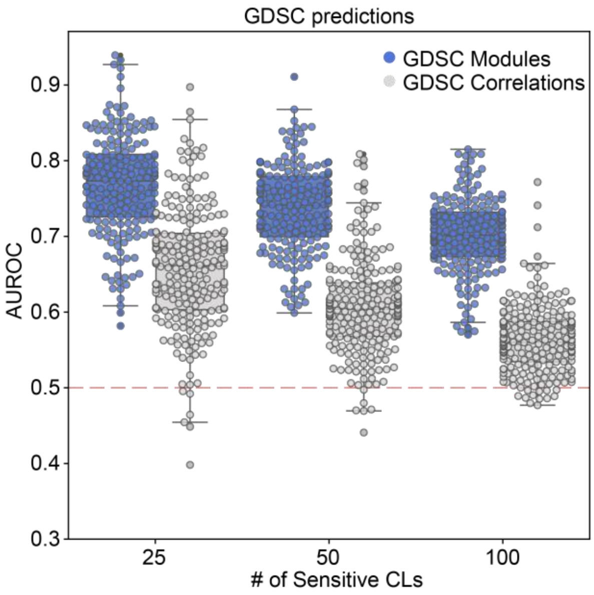


**Figure S13.** AUROC for the GDSC drug predictions using drug modules (blue) and significant drug-gene correlations (i.e. full signatures, gray).

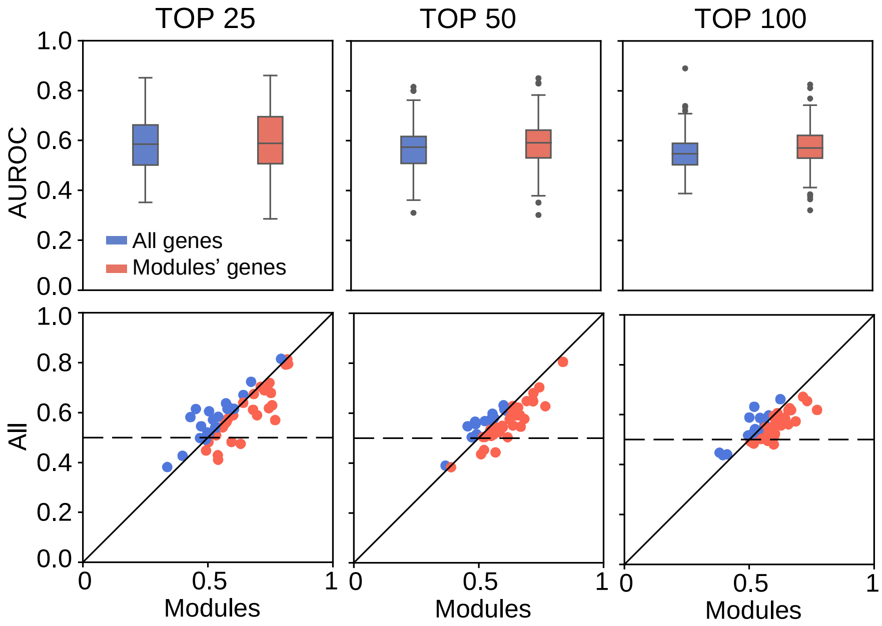


**Figure S14.** Performance of random forest predictors of drug sensitivity (a predictor was built for each drug; predictions for the top 25, 50 and 100 most sensitive cell lines are shown). (Top) Distribution of AUROC for the predictors using full gene expression profiles (blue), and module-specific profiles (red). (Bottom) A paired view of the AUROC values.

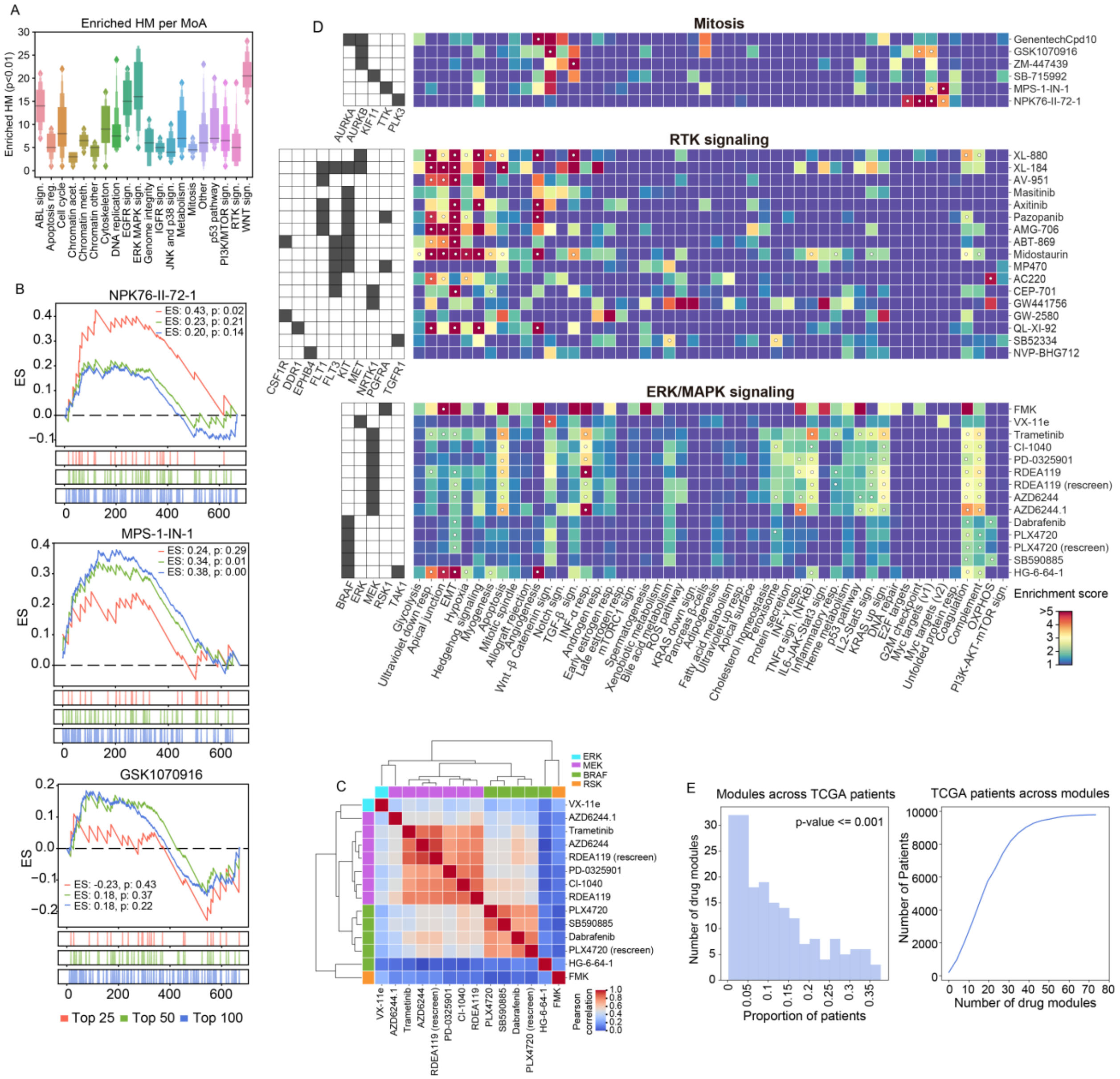

**Figure S15.** (A) Number of enriched Hallmarks (HM) (p-value < 0.01) across MoAs. (B) Myc gene expression enrichment in cell lines sensitive to NPK76-II-72-1 (top), MPS-1-IN-1 (middle) and GSK1070916 (bottom). (C) Hierarchical clustering between ERK/MAPK inhibitors drugs according their cell line sensitivity correlations. (D) Heatmap showing enrichment scores (odds ratio) of the positively correlated genes (full signatures) in the Hallmark gene set collection for 3 different drug classes: Mitosis (blue), RTK signaling (green) and ERK MAPK signaling. White dots denote significant enrichments (p-value < 0.05). (E) (Left) Number of enriched drug modules across the TCGA cohort. (Right) Cumulative distribution of the number of TCGA patients across the enriched drug modules.

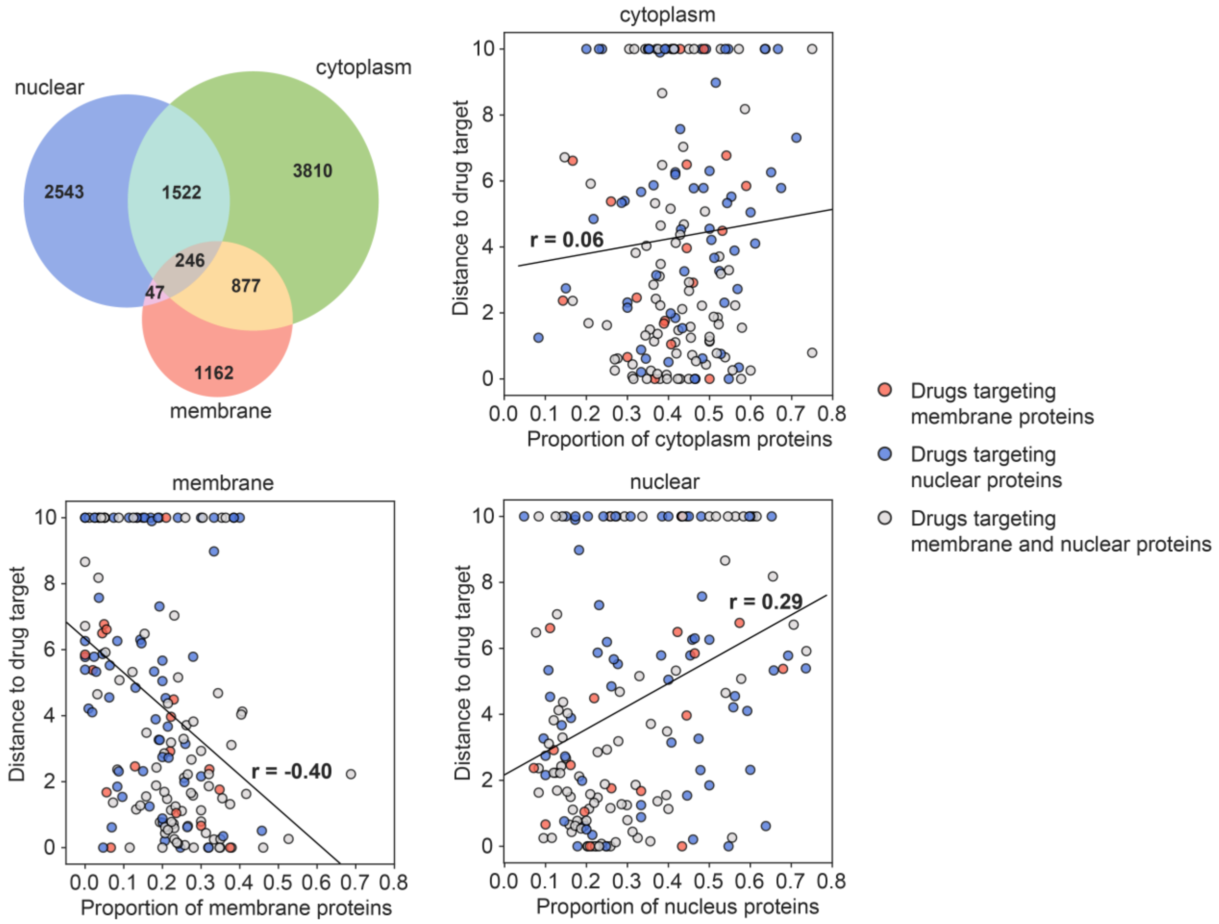


**Figure S16.** Correlation between the module distance to the drug target and the proportion of membrane, cytoplasm and nucleus proteins in positively correlated modules (PCMs; first module). The Venn diagram shows the proportion of proteins found in each cellular component category. In each correlation plot, blue dots correspond to drugs targeting a nucleus protein whereas red dots correspond to drugs targeting a membrane protein. In gray, we show drugs which target is found in both the nucleus and the membrane. Drugs with higher proportion of membrane proteins in their modules tend to have their target “nearby” the module (Spearman’s r = -0.40), while modules with more nucleus proteins tend to have more distal targets (r = 0.29).
